# Term Matrix: A novel Gene Ontology annotation quality control system based on ontology term co-annotation patterns

**DOI:** 10.1101/2020.04.21.045195

**Authors:** Valerie Wood, Seth Carbon, Midori A. Harris, Antonia Lock, Stacia R. Engel, David P. Hill, Kimberly Van Auken, Helen Attrill, Marc Feuermann, Pascale Gaudet, Ruth C. Lovering, Sylvain Poux, Kim M. Rutherford, Christopher J. Mungall

## Abstract

Biological processes are accomplished by the coordinated action of gene products. Gene products often participate in multiple processes, and can therefore be annotated to multiple Gene Ontology (GO) terms. Nevertheless, processes that are functionally, temporally, and/or spatially distant may have few gene products in common, and co-annotation to unrelated processes likely reflects errors in literature curation, ontology structure, or automated annotation pipelines. We have developed an annotation quality control workflow that uses rules based on mutually exclusive processes to detect annotation errors, based on and validated by case studies including the three we present here: fission yeast protein-coding gene annotations over time; annotations for cohesin complex subunits in human and model species; and annotations using a selected set of GO biological process terms in human and five model species. For each case study, we reviewed available GO annotations, identified pairs of biological processes which are unlikely to be correctly co-annotated to the same gene products (e.g., amino acid metabolism and cytokinesis), and traced erroneous annotations to their sources. To date we have generated 107 quality control rules, and corrected 289 manual annotations in eukaryotes and over 2.5 million automatically propagated annotations across all taxa.

## Introduction

The Gene Ontology (GO; http://geneontology.org) is the most widely adopted resource for systematic representation of gene product functions [1, 2, 3]. The core of the GO resource consists of two components: the Gene Ontology itself, and a set of annotations that use the ontology to describe gene products.

The ontology is a structured vocabulary that defines “terms” that represent biological structures or events, and the relations between them, in three interconnected branches: molecular function (MF; molecular-level activities of gene products), biological process (BP; larger-scale biological “programs” accomplished by multiple molecular activities), and cellular component (CC; the cellular locations in which a gene product performs a function). The ontology is structured as a graph, with class–subclass (*is_a*) relationships within each branch, and relationships of additional types (*part_of*, *regulates*, *occurs_in*, etc.) [4] within and between the three branches. Every GO term has a human-readable text definition, and a growing number have logical definitions that explicitly refer to terms in GO and other Open Biomedical Ontology (OBO) ontologies [2, 3, 5, 6]. (More formally, logical definitions use equivalence axioms expressed in OWL, the Web Ontology Language [7], to “specify necessary and sufficient conditions for class membership” for an ontology term.) Such definitions facilitate ontology structure maintenance and quality control.

GO annotations associate gene products with GO terms, with supporting evidence, a citation, additional metadata, and optional annotation extensions [8, 9]. (Note: annotations may use identifiers for genes as proxies for their products, and we use “genes” for simplicity in the remainder of this report.) The GO annotation corpus is widely used for a variety of genome-scale analyses, including broad characterization of whole genomes, interpretation of highthroughput transcriptomic and proteomic experiments, network analysis, and more [10, 11, 12, 13, 14]. In many cases, functional studies use subsets of the ontology (sometimes known as “GO Slims”), that exclude highly specific terms and take advantage of the fact that annotations are propagated over transitive relations (e.g., *is_a*, *part_of*) in the ontology.

Manual curation of primary literature describing experimental results supplies the most precise annotations. Experiment-based annotations are then propagated to genes from additional species by methods that include manual phylogeny-based transfer as well as computational methods using sequence models, orthology inferences, or keyword mappings [2, 15, 16, 17].

As a human endeavor, manual literature curation is imperfect, prone to errors in interpreting published experimental results or in choosing applicable ontology terms. In particular, the language used in publications is often less specific, or more prone to multiple interpretations, than the precisely defined ontology terms used in annotations. Furthermore, because manually curated annotations are widely propagated to support computed annotations, any inaccuracy in core manual annotation risks being transferred and amplified. Efficient ways to identify and correct errors are therefore highly valuable.

GO and model organism database (MOD) curators have developed a set of best practices to guide manual annotation, encompassing recommendations for interpreting various experimental results, selecting appropriate GO terms, applying evidence, and using annotation extensions [8, 18, 19]. Once created, annotations are subject to a series of automated quality control (QC) checks that flag errors for correction, such as incorrect term–evidence combinations, missing metadata, or file format problems [3]. Nevertheless, there is still ample scope for additional QC measures to improve the accuracy of the GO annotation corpus. Accordingly, we have developed a novel approach to annotation QC based on our observations of patterns of co-occurrence of different biological process terms used to annotate the same genes.

In biology, each gene may be involved in a wide variety of processes, and some have multiple functions; these are represented in GO as multiple annotations for a single gene. Due to spatial, functional or temporal constraints, however, certain combinations of functions or processes are not likely to be carried out by the same genes. We can therefore identify pairs of GO terms that are unlikely to be correctly annotated to the same genes, and thus provide a flag for potential mis-annotation. This work describes the development and initial implementation of a protocol that generates co-annotation QC rules from the identified pairs of GO terms to which the same gene should not be annotated, and then applies the rules in QC procedures to detect and correct annotation errors, and to prevent new occurrences of similar errors, thus yielding a higher quality annotation corpus.

## Methods

### Term Matrix annotation query tool

For each pair of GO terms analyzed, annotations were retrieved from the GO database by querying for gene products annotated to “Term1 AND Term2” directly or by transitivity (i.e. inferred over transitive relations in the ontology; by default, *is_a* and *part_of* are included). We developed a new tool, Term Matrix, which queries all pairwise combinations of a specified set of GO terms. Users can filter annotations by organism or annotated entity type (gene, protein, ncRNA, etc.) and can opt to include or exclude the *regulates* relations (*regulates*, *positively_regulates*, *negatively_regulates*) when traversing the GO graph to retrieve annotations inferred by transitivity. Results are displayed in a grid-based view (the “matrix”) that shows the number of gene products annotated to each pair of GO terms. Clicking the annotation count retrieves the annotation details for manual inspection.

Term Matrix uses the JavaScript D3 library. The code is released under the BSD 3-Clause “New” or “Revised” License, the same as the parent AmiGO application [20]. The tool works by querying the AmiGO Solr index, which uses precomputed graph closures that enable fast calculation of intersection counts for any term pair. Term Matrix is available directly [21], and accessible from the GO tools menu on the GO website [22].

### Annotation and ontology review

For pairs of GO terms with few co-annotated gene products, annotations and the cited sources were manually inspected to identify errors in manual literature curation or in mappings used to generate automated annotation. Where specific annotations appeared correct, the ontology was inspected for erroneous relationships.

We conducted several case studies, described below, to assess the effectiveness of our annotation validation process; the outcomes of the studies are discussed in the Results section.

### Fission yeast genome-wide annotation evaluation

*Schizosaccharomyces pombe* (fission yeast) annotations were evaluated at intervals (approximately biannually) over nine years. At each point, annotations to the then-current fission yeast GO slim [23] — a subset of GO biological process (BP) terms (usually about 40–50) selected to optimize coverage of informative cellular-level processes — were retrieved and assessed as described above. Before the Term Matrix tool became available, annotations to “Term1 AND Term2” combinations were retrieved by querying fission yeast annotations locally in PomBase (or its predecessor, GeneDB). Queries included annotations propagated over the *is_a*, *part_of*, and *regulates* relations in the go-basic version of the ontology [24]. Annotations were corrected, and queries re-run, iteratively.

### Cohesin complex annotation case study

We retrieved annotations to the GO cellular component term ‘cohesin complex’ (GO:0008278), a complex required for chromosome cohesion, combined with each of 35 GO BP terms, for all species in the GO database. Queries included the *is_a*, *part_of*, and *regulates* relations relations for BP ontology traversal.

### Cross-species GO subset case study

For cross-species analysis, we combined each of five selected terms [‘amino acid metabolism’ (GO:0006520),‘cytoplasmic translation’ (GO:0002181),‘ribosome biogenesis’ (GO:0042254), ‘tRNA metabolism’ (GO:0006399), and ‘DNA replication’ (GO:0006260)] with each term in a subset of 40 of the fission yeast GO slim BP terms. For six species (*S. pombe*, *Saccharomyces cerevisiae*, *Caenorhabditis elegans*, *Drosophila melanogaster*, *Mus musculus*, and *Homo sapiens*), we retrieved annotations in Term Matrix for each GO term combination, as described for fission yeast annotations. To avoid inclusion of genes involved in processes that have indirect effects, the *regulates* relations were not used for ontology traversal in this case study.

### Rule generation for annotation validation

Co-annotation QC rules were generated using the annotations retrieved in Term Matrix for the cross-species case study described above. After the correction of annotation errors, term pairs with no annotated gene products in common (mutually exclusive processes) were used to establish a set of rules capturing “Term1 is not usually co-annotated with Term2” statements. Rules are expressed in a simple tab-delimited text format, as described in Table 1. The set of rules that have been incorporated into GO’s annotation validation pipeline [3] is available at GO’s GitHub site [25], which also includes the runner code and additional tests and documentation [26]. Reports for the currently deployed GO release are available from 2018-08-09 onwards.

**Table 1:** Rule file format. Two mandatory columns contain the GO IDs for the pair of mutually exclusive terms, and remaining columns allow optional identifiers for exceptions to the rule (see “Allowable annotation overlaps” in the main text). For example, line 1 consists only of “GO:0006399 GO:0006457” in columns 1 and 2, and states that the GO terms ‘tRNA metabolic process’ (GO:00063996520) and ‘protein folding’ (GO:0006457) should not both be associated with a single gene. Column 3 may contain one or more pipe-separated IDs for GO terms that allow correct use of an otherwise mutually exclusive pair. In line 2, “GO:0006399 GO:0006310 GO:0045190” states that genes may be annotated to both ‘tRNA metabolic process’ (GO:0006399) and ‘DNA recombination’ (GO:0006310) only if they are annotated to ‘isotype switching’ (GO:0045190). Similarly, column 4 allows identifiers for individual gene products or for specific PANTHER families that cover entire orthologous groups, where annotation to both terms in a pair has been confirmed as accurate. In line 3, “GO:0002181 GO:0006605 WB:WBGene00006946” states that *C. elegans prx-10*, but not other genes, may be annotated to both ‘cytoplasmic translation’ (GO:0002181) and ‘protein targeting’ (GO:0006605) due to a tandem gene fusion in *C. elegans*.

| Term1 | Term2 | Excepted GO term | Excepted gene |
| --- | --- | --- | --- |
| GO:0006399 | GO:0006457 |  |  |
| GO:0006399 | GO:0006310 | GO:0045190 |  |
| GO:0002181 | GO:0006605 |  | WB:WBGene00006946 |

## Results

### Fission yeast genome-wide annotation evaluation

*Schizosaccharomyces pombe* has 5067 protein-coding genes, of which 4337 have annotation to a GO BP term more specific than the “root” term (‘biological process’, GO:0008150) [27]. The depth and breadth of fission yeast annotation make it an excellent system for studying co-occurrence of BP annotations. PomBase, the *S. pombe* MOD, maintains the fission yeast GO slim, a BP GO subset that classifies 99% of fission yeast protein-coding genes of known biological process into broad categories. Pairs of terms from the fission yeast GO slim with co-annotations were evaluated over time and visualized as described in the Methods. We thus identified term pairs that are rarely used to annotate genes in common, and then inspected the annotations individually.

We observed that the number of annotated genes shared by GO term pairs reflected biology: whereas large intersections between gene sets such as those annotated to ‘transcription’ (GO:0006351) and ‘chromatin organization’ (GO:0006325) are readily explained, biologically unrelated processes such as ‘tRNA metabolic process’ (GO:0006399) and ‘protein folding’ (GO:0006457) tended to yield few or no shared genes. Figure 1 illustrates the scale of annotation changes over time using annotation matrix “snapshots” based on data from 2012 and 2020 for 21 of the term pairs studied (before 2012, individual annotation error corrections were not systematically recorded). Individual annotation corrections derived from this analysis since 2012 are included in Supplementary Table S1.

**Figure 1:**
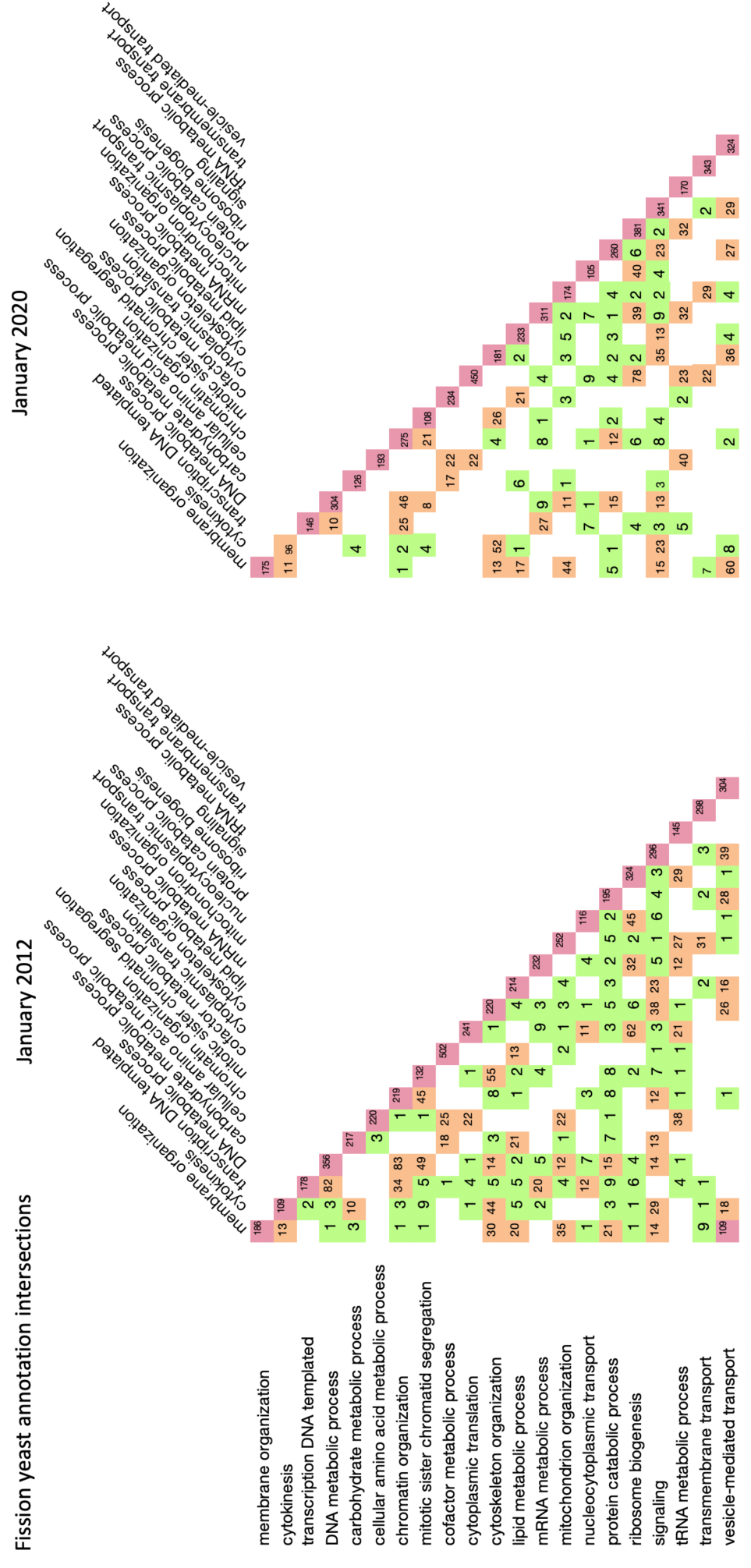
Annotation matrices showing fission yeast annotations for 21 selected GO term pairs in 2012 and 2020. Each row–column intersection off the diagonal shows the number of genes annotated to two different terms. Cells are colorcoded by number of co-annotated genes. Disputed phylogenetically-inferred annotations have been removed from the 2020 dataset.

### Annotation error types

Our work correcting fission yeast annotation errors led us to identify several classes of systematic error:

1. *Annotation of indirect effects:* In manual curation, incorrect annotations often arise when a phenotype is taken to mean that a missing/altered gene product normally participates directly in the process assessed, or measured, by the analysis, but is later shown to reflect a downstream effect of the mutation. For example, fission yeast Brr6 was originally thought to be involved in nucleocytoplasmic transport on the basis of the phenotype of the *S. cerevisiae* gene. Subsequent work showed that Brr6 in fact acts directly in nuclear envelope organization, and effects on nuclear transport lie downstream [27, 28]. In light of the most up-to-date knowledge, annotating Brr6 to ‘nucleocytoplasmic transport’ (GO:0006913) would be misleading. Likewise, perturbed DNA replication (GO:0006260) can indirectly lead to problems with chromosome segregation (GO:0007059), due to the presence of DNA structures that cannot be separated (e.g. [29]. A chromosome segregation phenotype alone therefore does not suffice to confidently annotate a gene product as involved in chromosome segregation. In more extreme cases, downstream effects of mutations can sometimes lead to erroneous annotation of genes that do not normally influence a process, even indirectly. Cell cycle arrest phenotypes often give rise to this type of error, because the arrest may result from mutations that cause problems that a functioning checkpoint can detect but not correct. For example, decreased expression of the ribosome processing protein SLBP results in slowed cell growth and an accumulation of cells in S phase. From these phenotypes, SLBP was erroneously annotated to terms ‘DNA replication’ (GO:0006260) and ‘cell cycle phase transition’ (GO:0044770), despite playing no role in either process in a normally functioning cell. Indirect effects are frequently seen, and most at risk of misinterpretation, in high-throughput datasets where candidate genes may be annotated without data from follow-up validation experiments.
2. *Term interpretation and usage:* Errors in manual curation can arise from misinterpretation of experimental results or the meaning of a GO term. For example, we found 12 examples of genes annotated to ‘transmembrane transport’ (GO:0055085) where ‘nucleocytoplasmic transport’ (GO:0006913) would instead be correct, because during nucleocytoplasmic transport the lipid bilayer is not traversed. Occasionally enzyme activities are misinterpreted; e.g. *S. cerevisiae KTI1*, an oxidoreductase involved in tRNA wobble uridine modification, was annotated to ‘electron transfer activity’ (GO:0009055). This molecular function term specifically represents the action of an electron acceptor and electron donor in an electron transport chain, and is linked directly to the biological process ‘electron transport chain’ (GO:0022900); it is more specific than the oxidoreductase activity of *KTI1*.
3. *Mappings:* Because manually assigned experimental annotations provide the main source of data to create automated annotations, all types of annotation error described above can result in the incorrect association of GO terms to InterPro signatures (InterPro2GO mapping) [2, 30] or UniProt keywords [2, 31]. In addition, other error types specifically affect annotation derived from automated mappings. First, irrelevant terms can be propagated, either via matches to domains found in proteins from species in which a process, activity, or cellular location does not exist, or via transfer of a very specific GO term instead of a less precise, but more broadly applicable, GO term (in our study, 13 families had mappings which were only true for a subset of entries; these were excluded from the annotation error count). Second, mappings derived from protein family membership can be affected by false positive family assignments.
4. *Ontology structure:* Incorrect paths in the ontology can cause erroneous inferences to “ancestor” terms from correct annotations to “descendant” terms. For example, the parent ‘citrulline biosynthetic process’ (GO:0019240) has been removed from ‘protein citrullination’ (GO:0018101), because protein citrullination describes the modification of an amino acid residue in a protein into citrulline, not the synthesis of free citrulline.
5. *Advances in biology:* Older findings can be supplanted by new knowledge, especially paradigm shifts in biology. For example, the Elongator complex was long thought to act as a histone acetyltransferase (GO:0004402), based on assays that have since been shown to be artefacts (e.g. see [32]). Instead, the Elongator complex is actually involved in tRNA modification (GO:0006400), and all observed phenotypes can be attributed to this role [33, 34]. Annotations need to be adjusted to reflect this new knowledge.

### Allowable annotation overlaps

After correcting errors in manual annotations, mappings used for automated annotation, and ontology relationships, many GO term pairs had no annotated gene products in common. The exceptions all fall into one or more of the following types:

1. Annotation to a term that is a descendant of both assessed terms. For example, ‘pentose-phosphate shunt’ (GO:0006098) has paths to both ‘nucleotide metabolism’ (GO:0009117) and ‘carbohydrate derivative metabolism’ (GO:1901135).
2. Gene products involved in regulatory pathways upstream of both processes, usually signalling pathways (GO:0007165), gene expression (GO:0010467), or protein catabolism (GO:0030163).
3. Multifunctional gene products, tandem fusions and moonlighting proteins. For example, *S. pombe* Noc3 functions in both DNA replication (GO:0006260) and rRNA processing (GO:0006364).

### Interspecies case studies

#### Cohesin complex

We next conducted two case studies to investigate whether the utility of coannotation analysis for annotation QC would hold for species other than the well-annotated fission yeast. In the first, we examined co-annotations to the GO cellular component term ‘cohesin complex’ (GO:0008278) with each of 35 selected GO BP terms.

Erroneous annotations fell into the same categories identified for fission yeast annotations. Figure 2 shows co-annotation counts before and after corrections, and a breakdown of annotation errors by type and database. Across multiple MODs plus UniProt, 35 experimental annotations were deleted (listed in Supplementary Table S2). Finally, one InterPro2GO mapping affecting over 7000 computationally inferred annotations was removed (see Supplementary Table S3).

**Figure 2:**
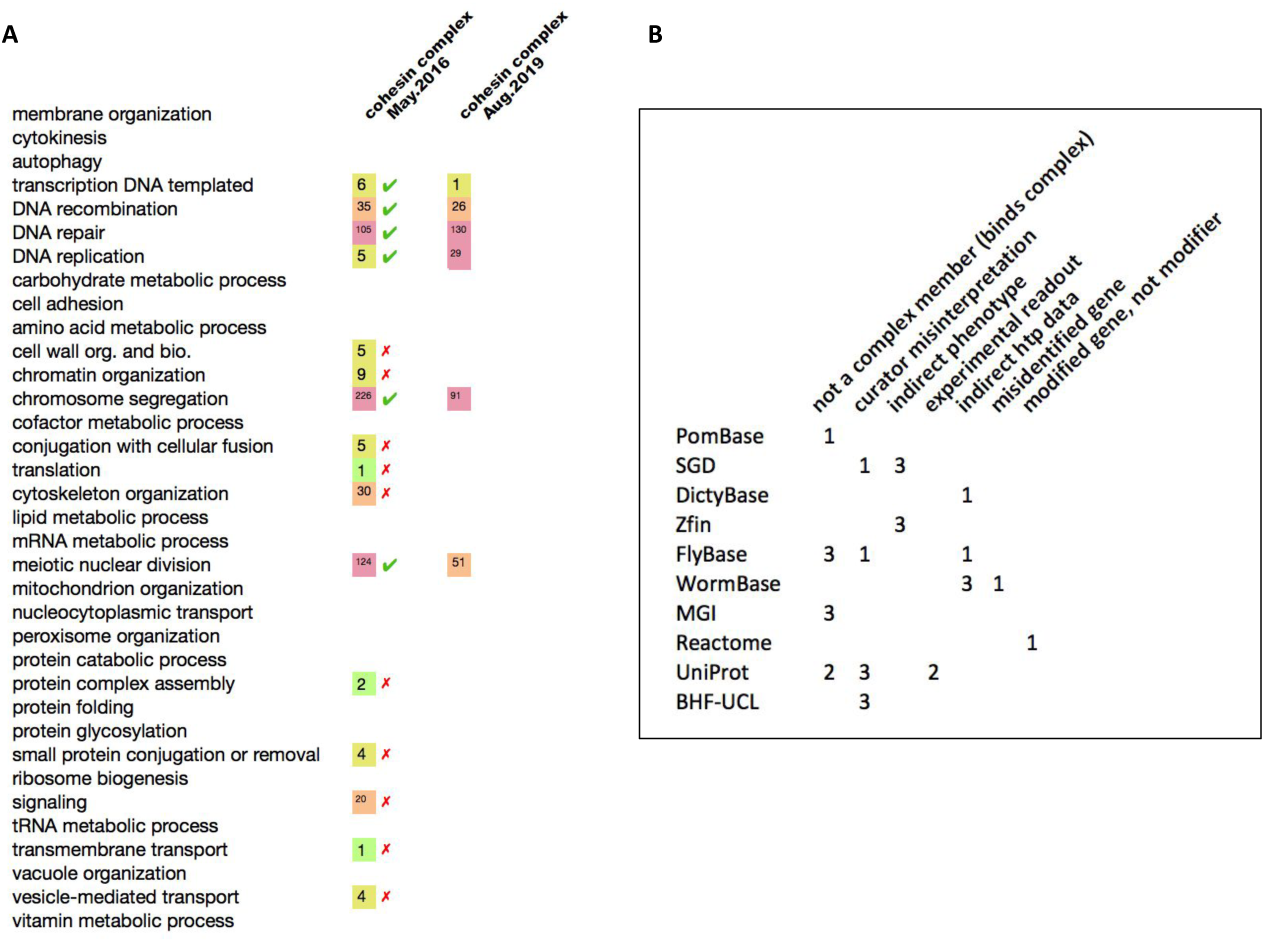
**A.** For each of 35 GO BP subset terms, the cumulative number of genes in all organisms annotated to both the BP term and the cellular component term ‘cohesin complex’ (GO:0008278) is shown for May 2016 and August 2019. **B.** For each database, the table shows the number of annotation errors of each type identified and corrected.

#### GO biological process subset

In the second cross-species study, we narrowed the number of species to six (fission yeast, budding yeast, worm, fly, mouse, and human; see Methods), but broadened GO coverage to pairwise combinations of five core cellular level biological processes [‘amino acid metabolism’ (GO:0006520), ‘cytoplasmic translation’ (GO:0002181), ‘ribosome biogenesis’ (GO:0042254), ‘tRNA metabolism’ (GO:0006399), and ‘DNA replication’ (GO:0006260)] against a set of 40 core cellular level biological process GO terms (Supplementary Table S4). As a result, 182 manual annotations were corrected or removed; over two million annotations were addressed by correcting 19 ontology paths; over 380,000 inferred annotations across all species based on 54 InterPro2GO mappings (based on the family size in InterPro version 77) were corrected; and over 1800 annotations for key GO species phylogenetically inferred using the PAINT annotation transfer system [16] from 14 PANTHER protein families were corrected (see Supplementary Table S3 for InterPro2GO and PANTHER mapping corrections). Finally, two UniRule [35] and 41 UniProt keyword mappings were revised or deleted (Supplementary Table S5). As for the fission yeast study, manual annotation corrections made for the BP slim analysis are included in Supplementary Table S1. Supplementary Table S6 lists ontology corrections and numbers of affected annotations.

#### A workflow for annotation quality control

Following successful detection and correction of annotation errors in our case studies, we have developed shared co-annotation rules that form the basis of a pipeline for annotation QC. The “Matrix QC” workflow is a multi-step, ongoing and iterative process, summarized in Figure 3:

1. A set of GO term identifiers is used as input for the Term Matrix tool to provide visualization and access to genes with annotations shared between pairs of GO terms (annotation intersections). Early iterations use selected GO terms, and use the Term Matrix option that excludes *regulates* relations when traversing ontology paths (i.e. gene products annotated to a term that is connected to one or both queries via the *regulates* relation will not appear in the intersection set). Annotation outliers, defined as intersecting sets with low numbers of annotated gene products, are critically inspected for validity. Annotation errors are identified and corrected, usually by assessing the original experimental data. As part of establishing the Matrix QC workflow, we corrected 289 manual annotations, 55 InterPro2GO mappings, 14 PAINT propagation errors from PANTHER families, two UniRule mappings, 41 UniProtKW mappings and 19 ontology paths, as summarized in Table 2. Annotation intersections which yield empty sets can be used to generate co-annotation QC rules of the form “genes annotated to process A are not usually annotated to process B”.
2. New and existing annotations that violate annotation co-annotation QC rules are reported to contributing databases via standard GO Consortium QC pipelines.
3. Upon reviewing reported errors, the contributing database may either make corrections or provide evidence that validates annotations. For valid annotations in intersections, the co-annotation QC rules are extended to include additional specifications that will allow only valid annotations to pass (exceptions may be specified at the level of species, protein families, or individual gene products). Rules can also be modified to account for new biological knowledge, for example by allowing co-annotation of specific sub-processes, or where the annotated gene product matches additional criteria such as being annotated to a particular molecular function or cellular component.

**Figure 3:**
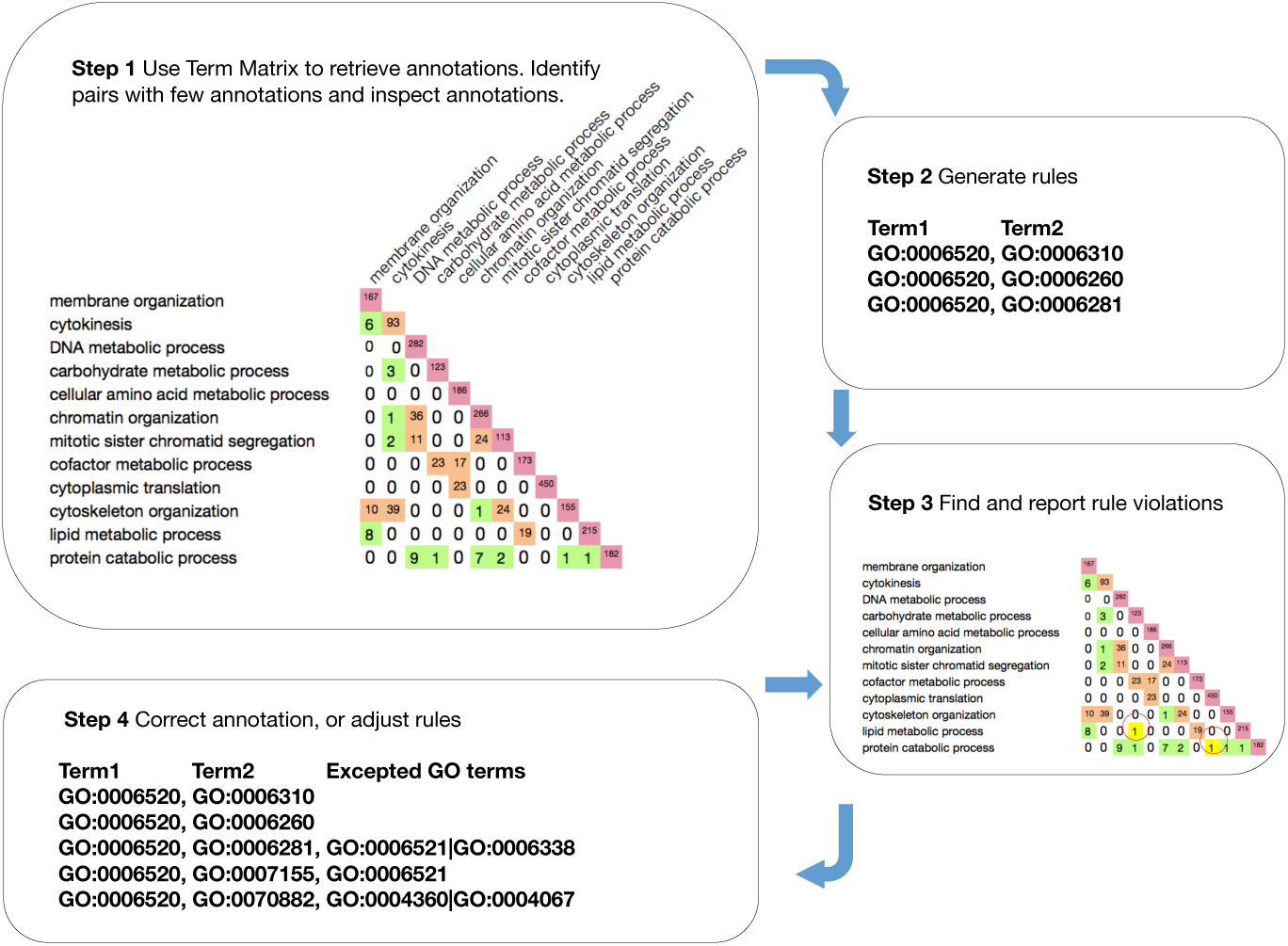
Intersection-based annotation quality control workflow. Step 1: Term Matrix retrieves annotations shared between pairs of GO terms. For term pairs with few annotations, both annotations and ontology are inspected, and errors corrected. Step 2: Based on known biology, create co-annotation QC rules that disallow simultaneous annotation to term pairs (“NO OVERLAP” between annotation sets for the indicated terms). Step 3: Re-run Term Matrix to find annotations that violate the rules; report to contributing databases for validation. Step 4: Correct annotation errors, or amend rules to allow specific biologically valid exceptions.

**Table 2:** Error types. Number of different errors of each type found in annotations and the ontology structure, and the number of annotations affected. ND: not determined.

| Incorrect mappings | Occurrences | Entries affected |
| --- | --- | --- |
| UniProt keyword to GO mapping | 41 | ND |
| UniPathway to GO mapping | 2 | ND |
| InterPro to GO mapping | 55 | >380,000 |
| PAINT annotation | 14 | 1818 |
| **Ontology corrections (incorrect parent)** |  |  |
| Affecting all annotations | 19 | >2,000,000 |
| **Non-systematic manual annotation error** |  | 289 |

## Discussion

### Using co-annotation for quality control

Biological data can sometimes be subject to variable interpretation, and the state of biological knowledge is constantly changing, posing challenges for the accurate and up-to-date characterization and curation of genes and their products. We have developed a QC pipeline for GO BP annotation based on observed patterns of co-occurrence of GO terms used to annotate the same genes. Annotation to both of a selected pair of GO terms, designated “annotation intersection”, should occur only where the processes actually overlap, genes are shown to have multiple functions, or the same function is used in more than one process. We have corrected numerous errors in annotations and ontology relationships, generated rules describing annotation intersections expected to be null, and incorporated the rules into an iterative QC pipeline. The new system provides for the detection and correction of existing annotation errors, as well as prevention of similar errors entering the GO annotation corpus.

Our work demonstrates that the inspection of annotations co-annotated to multiple processes can identify annotation outliers, systematic mapping errors, and ontology problems for validation or correction. The incremental creation of co-annotation QC rules covering all annotation space will create a robust mechanism for the validation and improvement of the annotation corpus over time, because potential errors will be identified, and the flagged annotations validated or corrected upon submission.

### Propagating error correction

Errors in ontology relationships and in mappings between ontology terms and other classification systems such as InterPro can introduce systematic errors in annotation datasets, as often every gene product annotated to a misplaced term, or associated with a particular domain or keyword, is affected. Systematic errors may also originate from experimentally supported annotations produced by curators, because these manually curated data are then used to develop mappings between GO and protein families, and in phylogenetic propagation. For example, experimental annotations assigned by MOD and UniProt curators are used to establish InterPro2GO mappings, and routinely propagated to orthologs across thousands of species by Ensembl Compara pipelines [36] and PAINT. Furthermore, widely propagated annotation errors affect common uses of GO data; for example, misannotation of many genes to the same term in a given species can obscure enrichments. Correcting these errors makes a correspondingly wide-reaching improvement in the GO annotation corpus, affecting hundreds or even thousands of annotations, as well as in analyses that use annotation datasets.

### Future directions

In the present study, we created co-annotation QC rules for pairwise term combinations involving five GO BP terms that already had low numbers of annotations in intersections. We aim to extend the rules to cover more GO term pairs, and to accommodate all experimentally verified annotations found in intersections using increasingly specific rule exceptions. Although rule construction and the accompanying error correction procedure is time-consuming — because large numbers of potential violations need to be traced back to the original publication and evaluated, and the reasons for the apparent violation are often obscure — maintenance overhead is low once rules are established. Adapting co-annotation QC rules and exceptions to accommodate new biology takes comparatively little effort, and provides annotation quality benefits indefinitely.

Next, we will extend co-annotation QC rules to the MF and CC branches of GO, adding rules for pairwise combinations of terms within the MF and CC branches, and for pairs of terms from different branches. For example, a rule could identify cytosolic (CC) proteins that are annotated to DNA recombination (BP), or that a DNA-binding transcription factor activity (MF) is not a general transcription initiation factor activity (MF). We will also explore applications of co-annotation QC rules beyond error detection in existing annotations. For example, machine learning function prediction exercises may use our QC rules to constrain predictions such that annotations that would violate rules are excluded.

We also anticipate that combining Term Matrix-based QC with novel GO annotation protocols will yield synergistic benefits. Our results to date indicate that, due to downstream or pleiotropic effects, it is difficult to assign a direct role in a biological process to a gene product from a mutant phenotype without additional information. Additionally, it can often be challenging to discern when a gene product is directly involved in a biological process as opposed to having an impact on a process by perturbing an upstream process. Two recent innovations in GO annotation show great promise for minimizing such errors. First, the introduction of new relations to describe how a gene product is connected to a term (*involved_in*, *acts_upstream_of*, etc.), will allow curators to capture indirect annotations explicitly, and simultaneously provide a mechanism to filter when a set of direct annotations is desired. Second, the new gene product–GO term relations form part of the Gene Ontology Causal Activity Modeling (GO-CAM) system, which uses OWL to represent how the molecular activities of gene products interconnect to carry out, and regulate, biological processes.

### Conclusions

We envisage that the co-annotation rule-based QC procedure will help direct researchers to outstanding questions in molecular or cellular biology: annotations that appear to violate rules may indicate areas where available experimental results are inconsistent, requiring further experiments to resolve discrepancies, but some may identify interesting areas of biology where evolution has co-opted a single gene product for more than one task. Our work has built a co-annotation QC system into GO procedures that can readily be more widely implemented, thereby enabling curators and researchers to distinguish between new annotations that provide additional support for known biology and those that reflect novel, previously unreported connections between divergent processes. The coannotation QC pipeline thus enhances GO not only by detecting and preventing annotation errors, but by highlighting advances in our understanding of biology.

## Acknowledgements

We thank Peter D’Eustachio for Reactome updates and the InterPro group for InterPro2GO mapping updates. We thank Nomi Harris for constructive comments on the manuscript. We also thank the many biocurators, editors, and other members of the GO Consortium who have contributed GO annotations and to the development of the Gene Ontology.

## Data accessibility

The GO ontology and annotation datasets are freely available from the Gene Ontology website (see the main downloads page [38]). All other data supporting this article have been uploaded as part of the supplementary material.

## Authors’ contributions

VW conceived the project and wrote the initial draft; SC and CJM developed Term Matrix; KMR provided bioinformatic support for the fission yeast case study; VW, AL, SRE, DPH, KVA, HA, and RCL corrected annotation errors identified in the study; MAH made extensive text revisions, and prepared the manuscript for submission; DPH, KVA, and PG corrected ontology errors; SP and MF provided SPKW mapping updates; MF and PG provided PAINT propagation updates. All authors contributed to the discussion of ideas and manuscript revisions, and read and approved the final manuscript.

## Competing interests

The authors declare no competing interests.

## Funding

VW, AL, MAH, and KMR are supported by the Wellcome Trust (grant no. 104967/Z/14/Z). The GO resource (SC, SRE, DPH, KVA, PG, CJM) is supported by the National Human Genome Research Institute (NHGRI) (grant no. U41 HG002273). SRE is also funded by the NHGRI via the *Saccharomyces* Genome Database (grant no. U41 HG001315) and the Alliance of Genome Resources (grant no. U24 HG010859). KVA is also funded via WormBase, which is supported by the NHGRI (grant no. U24 HG002223), the UK Medical Research Council (grant no. MR/S000453/1) and the UK Biotechnology and Biological Sciences Research Council (grant no. BB/P024602/1). HA is funded by the UK Medical Research Council (grant no. MR/N030117/1).

## Notes

### Competing Interest Statement

The authors have declared no competing interest.

